# An overview of GabRat edge disruption and its new extensions for unbiased quantification of disruptive camouflaging patterns using randomization technique

**DOI:** 10.1101/2024.02.27.582325

**Authors:** Masahiko Tanahashi, Min-Chen Lin, Chung-Ping Lin

## Abstract

Disruptive colorations are camouflaging patterns that use contrasting colorations to interrupt the continuously of object’s edge and disturb the observer’s visual recognition. The GabRat method has been introduced and widely used to quantify the strength of edge disruption. The original GabRat method requires a composite image where a target object is placed on a particular background. It computes the intensities of ’frequency components’ parallel and perpendicular to the edge direction at each edge point using Gabor filters, and summarize the ratios of these two intensities around the perimeter of the shape. However, we found that the original GabRat method has an issue which produces false signals and biases to overestimating the GabRat value depending on the edge angle. Here, we introduce GabRat-R, which can diminish that angle dependency using Gabor filters with randomized base angles. Additionally, we developed GabRat-RR, which iteratively places a target object on a background with random positions and rotation angles to average the effects of the heterogeneity and anisotropy of background. Compared with the original GabRat, our GabRat-R and GabRat-RR programs run more efficiently using multithreading techniques. GabRat-R and GabRat-RR were freely available in *Natsumushi* 2.0 software of the authors’ website.

## Introduction

Detection of continuous edge is known to be one of the most fundamental process of visual recognition of target objects in many animals such as human [1][2][3], mammals [4], birds [5], fishes [6] and insects [7]. Visual animal predators predominantly use preys’ body outlines to find their preys [8]. To avoid attack, the preys can evolve camouflage to obscure themselves as the first line of defense against the predators [9]. One form of camouflage coloration is the disruptive marginal patterns in which colors intersect the body edges of the preys. Such colorations are thought to have a disruptive effect on the visual continuity of the outline of organisms, and thus, they can serve as camouflage to conceal the real object’s body boundary [10]. Here is an example of a disruptive camouflage, in which contrasting coloration patterns are in contact with the edge of object’s outline (Fig 1). When the object is put on a white background, the outline of the object is less disruptive because the contrast between the object’s colorations and the background (blue vs. white) is higher than the contrast within the object (blue vs. black) (Fig 1a). However, when the object is on a green background, the contrast between the object and the background (blue and green) becomes lower than the contrast within the object (blue vs. black) (Fig 1b). That produces an illusionary outline (red line), and consequently, the true edge of the object is obscured (Fig 1c). Various methods have been developed to quantify the degree of edge disruption of an object [11][12][13][14][15][16]. Some quantitative methods use simple mathematical operations, such as Gabor filters, to provide relatively objective and versatile evaluation of edge disruption [16], whilst other methods rely more on the methods of computer visions, such as Canny edge detector [11] and SIFT feature detection [16], which eventually require the thresholding processes that are sensitive to the artificially-defined parameters. In this study, we first present an introduction of GabRat edge disruption, which was originally developed by Troscianko *et al*. (2017)[16]. Secondly, we identify several issues associated with the calculation of GabRat. Finally, two new methods, namely, GabRat-R and GabRat-RR are proposed to solve those issues and extend the application of the original GabRat.

**Fig. 1.**
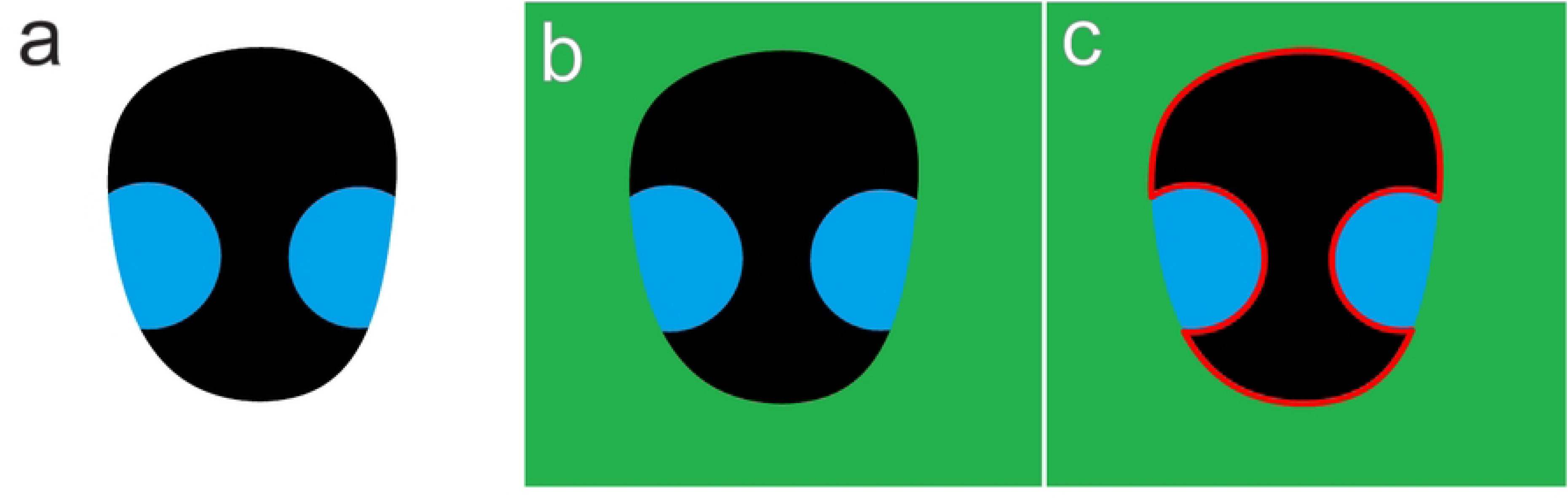
Basic concept of edge disruption effect. (a) A virtual insect image that have black body surface with two blue colorations crossing on the edge. (b) The virtual insect is put on a green background. (c) In this case, visual recognition of the object is disrupted by the blue colorations that create more contrastive false boundaries (red line).

### Overview of basic GabRat calculation

GabRat is a method designed to quantify the effect of edge disruption when a target object is placed on a specific background image [16]. Strength of edge disruption can be defined as the relative intensity of contrastive local patterns perpendicular to the edge of the object, because such patterns often disturb the visual recognition of true object’s outlines. GabRat uses Gabor filters [17], which is generally utilized to determine the intensity of spatial frequency contents in specific directions. Fig 2 shows examples of extracting spatial frequency contents in vertical and horizontal directions, using the simplest 3 x 3 Gabor-like filters. Before the filter operation, a digital color image (Fig 2a) is converted to a monochrome image or separated into RGB channels, so that each pixel of the image has a single intensity (brightness) value of 0.0 to 1.0. To calculate the vertical energy at a specific pixel (= *E_i_*, *_j_*), the intensities of an upper-left neighborhood pixel (= *P_i_*_–1,_ *_j_*_–1_) is multiplied by the value of the filter matrix at the corresponding location (= 1), the upper neighborhood pixel (= *P_i_*, *_j_*_–1_) is multiplied by zero, the same applies hereafter, and the results are summed up as follows:

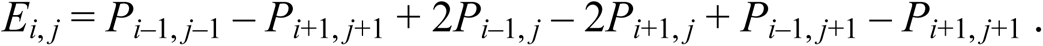

**Fig. 2.**
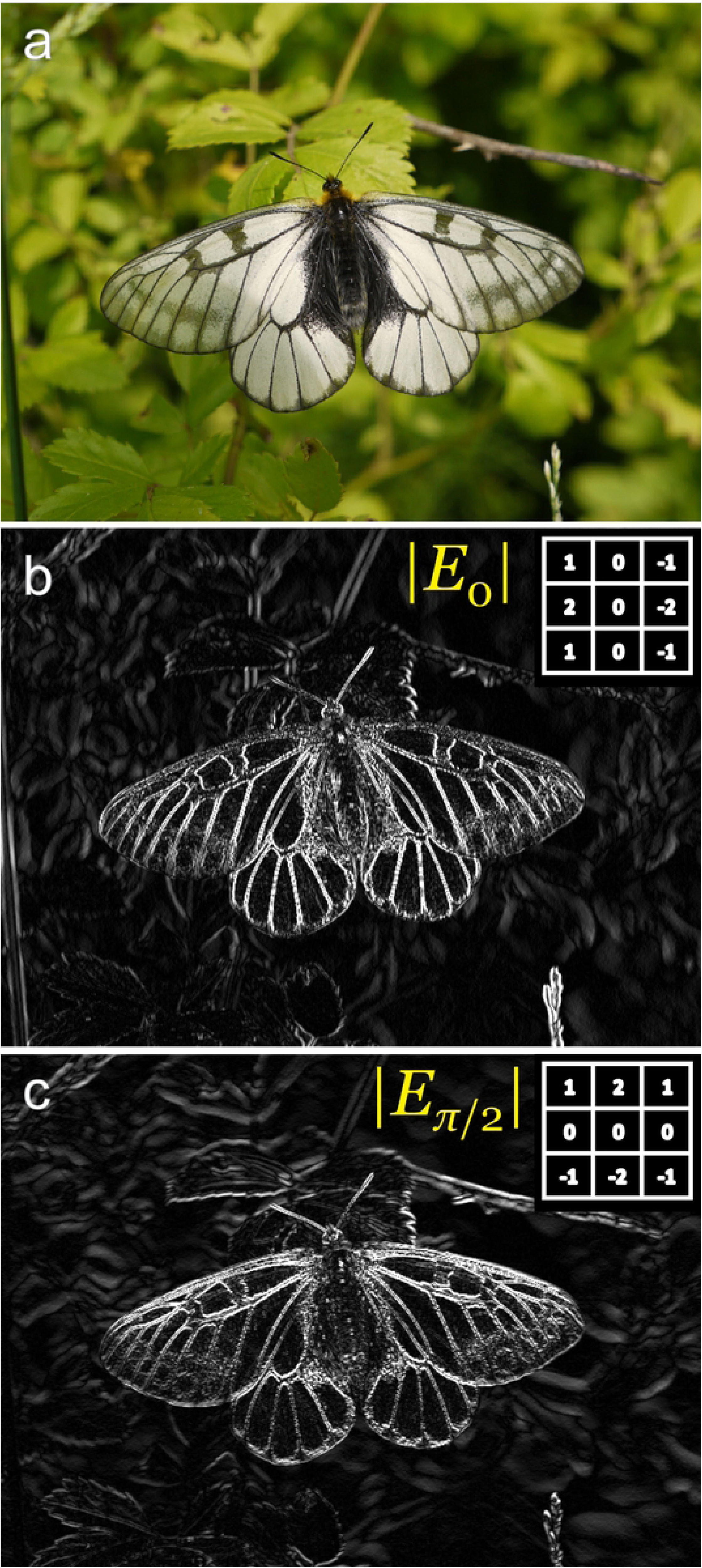
Examples of the simplest, 3 × 3 Gabor filter matrices and their outputs. (a) A sample photograph of the glacial Apollo butterfly, *Parnassius glacialis*. (b) Absolute vertical edge energy (*θ* = 0°) and (c) absolute horizontal edge energy (*θ* = 90°).

Since the positives and negatives of the filter matrix elements are reversed across the vertical axis, perpendicular patterns (e.g. horizontal zebra patterns) are canceled in the matrix calculation. As a result, the vertical Gabor filter selectively passes the spatial frequency contents with vertical patterns (Fig 2b). Similarly, horizontal energy of the image at each location can be calculated using the horizontal Gabor filter (Fig 2c).

GabRat calculation uses a set of Gabor kernel filters that is determined by four parameters: kernel size (*σ*), aspect ratio (*γ*), frequency (*Fx*) and number of filter angles (*nAngles*). Of those, *σ* needs to be determined for each analysis, depending on the pixel size of the target object, so that the kernel filter sufficiently covers the edge boundary and its surrounding area (the size of the filter matrix, *k × k*, is determined as *k =* 6*σ* + 1). The aspect ratio *γ* is basically fixed to 1.0. Changes of *Fx* alter the frequency of the zebra pattern of Gabor filter (S1a Fig); however, it is not recommended to use a large *Fx* value (>2.0) because the result becomes exclusively sensitive to the specific recurrent pattern with that frequency. In this study, we only tested *Fx* = 2.0, following the previous studies [16][18].

The *nAngles* parameter is the essence of GabRat calculation, and also the root of all issues identified in the next section. In GabRat calculation, typically, four Gabor kernel filters of different rotation angles (*nAngles =* 4; 0°, 45°, 90° and 135°) (S1b Fig) have been used to determine ’edge energy’ of specific direction around each edge point [16][18]. GabRat first determines the ’edge angle’ on each pixel that constitutes the edge outline. To do this, it temporally generates a binary (black-and-white) image, in which black (= 0) represents the background and white (= 1) represents the area of the foreground object (Fig 3a). GabRat picks every edge pixel and calculates the edge energies of four different directions on the binary image using Gabor kernel filters corresponding to those angles. Hereafter, the angle at which the absolute energy (i.e. absolute value of the edge energy) becomes highest is called the parallel angle (*θ_par_*). It should be noticed that *θ_par_* is not always exactly parallel with the tangential line of the true edge. In other words, the continuous edge angle is discretized into four angles (0°, 45°, 90° and 135°) by this operation. The orthogonal angle (*θ_ort_*) is simply assigned to be perpendicular to *θ_par_*, i.e.: *θ_ort_ =* (*θ_par_* + 90°) mod 180°. Next, the same GabRat filters are applied to the real (grayscale) image in order to calculate four absolute energies: |*E_0_*|, |*E_45_*|, |*E_90_*| and |*E_135_*| at the corresponding edge point (Fig 3b). Of those, the parallel energy |*E_par_*| and orthogonal energy |*E_ort_*| are chosen according to the pre-determined edge angles *θ_par_* and *θ_ort_*, respectively. Notice that when the real image has some disruptive patterns at that location, |*E_par_*| does not always indicate the maximum value of the four absolute energies (Fig 3b, position II). Finally, GabRat value on that edge point is defined as: *GabRat* = |*E_ort_*| / (|*E_par_*| + |*E_ort_*|). According to this definition, GabRat value exceeding 0.5 means that the orthogonal energy is larger than the parallel energy at that specific location, therefore, indicates that the false edge is more contrastive than the true edge. In most analysis, GabRat values are summed up for all edge pixels on the object’s contour and the mean GabRat value is used to represent the level of edge disruption of the target object against the specific background [16].

**Fig. 3.**
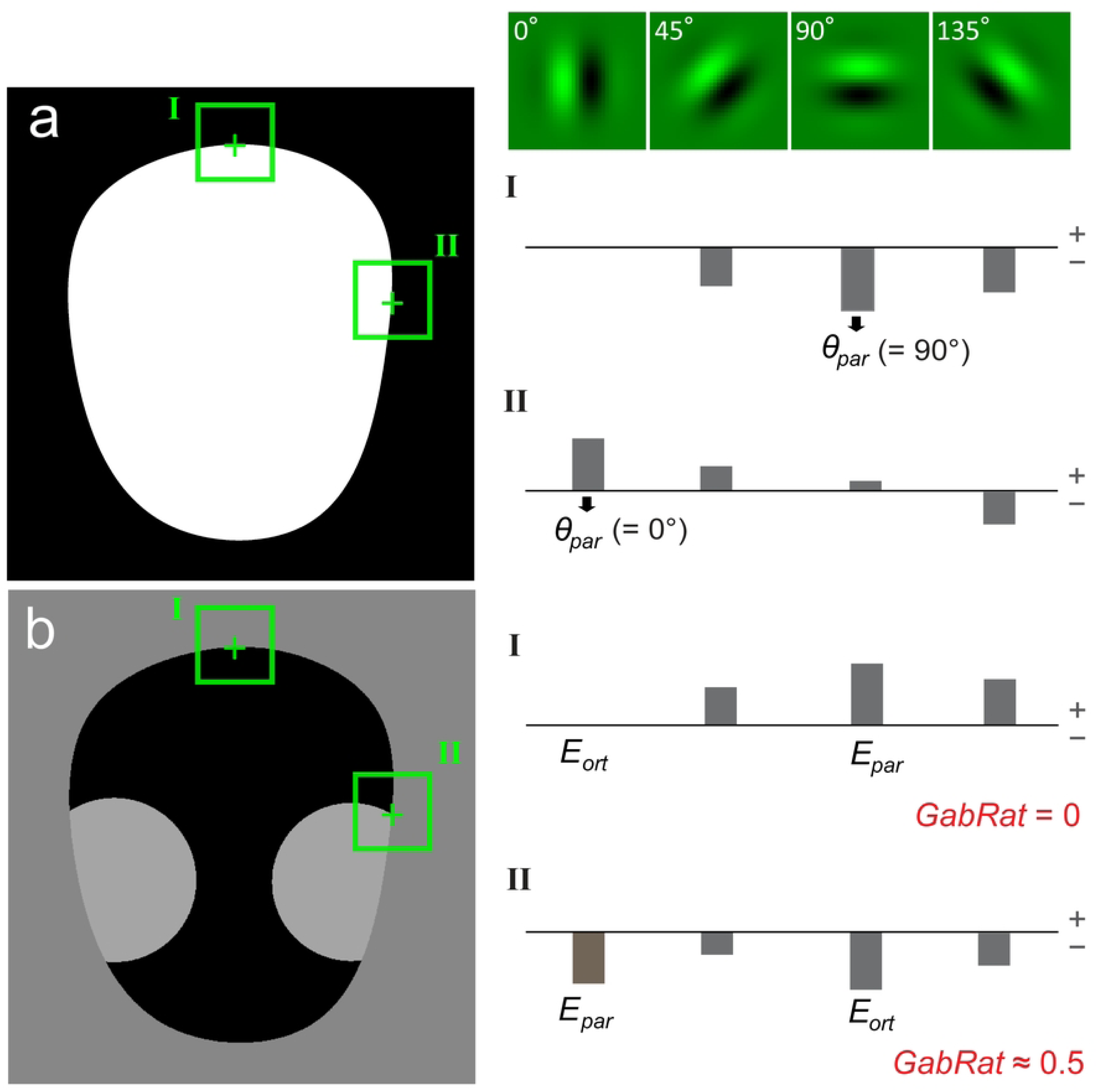
The principle of GabRat calculation. The model image is same as Fig. 1a. (a) At the first step, edge energies of four different angles (*nAngles = 4*) are calculated on a binary (black and white) image along the object edge using Gabor filters. The filter angle that gives the maximum absolute edge energy (*θ_par_*) is referred to as the ’edge angle’ at each location. (b) Next, edge energies of four different angles are calculated on a gray scale image in the same way. Of those, edge energy at the angle of *θ_par_* is chosen to be the parallel energy (*E_par_*). Orthogonal energy (*E_ort_*) is simply defined as the edge energy that is perpendicular to *θ_par_*. A local *GabRat* value is calculated as *GabRat* = |*E_par_*| / (|*E_par_*| + |*E_ort_*|) on each point.

The original GabRat was implemented as a plugin function of mica Toolbox [15] of ImageJ software [19][20] and the Java source code of the original GabRat program was deposited to the public code depository website, GitHub (https://github.com/troscianko/micaToolbox) under the GPL-3.0 license.

### Issues in GabRat calculation

Although GabRat value is a useful index to quantify the effect of edge disruption, we noticed that the original GabRat calculation has an intrinsic issue, which causes the angle-dependent false signal pattern. Fig 4 shows this false signal pattern using the simplest foreground and background models. When a white circular object is put on a black background (Fig 4a), the strength of edge disruption is expected to be zero on every edge point, since the object has no apparent false edge. However, the original GabRat calculation (*σ* = 6, *nAngles* = 4) exhibits an angle-dependent periodic signal pattern, the intensity of which ranges from 0.000 to 0.188 (mean = 0.085) (Fig 4b and Table 1). This type of false signal pattern is more pronounced with shapes that include straight lines, especially when the angle of the straight line is distant from a multiple of *π*/4 (Fig 4d, e). Moreover, the intensity of the false signal is dependent on the rotation of the entire image. In Figure 5, we first prepared a composite image in which an insect model with edge disruptive patterns was placed on a background, then rotated the entire image by *π*/8 (= 22.5°). Although those two composite images were identical except for their placement within the computer screen, the mean GabRat values varied from 0.030 to 0.173, which is mainly caused by the false signals on the edges with straight lines (Fig 5).

**Fig. 4.**
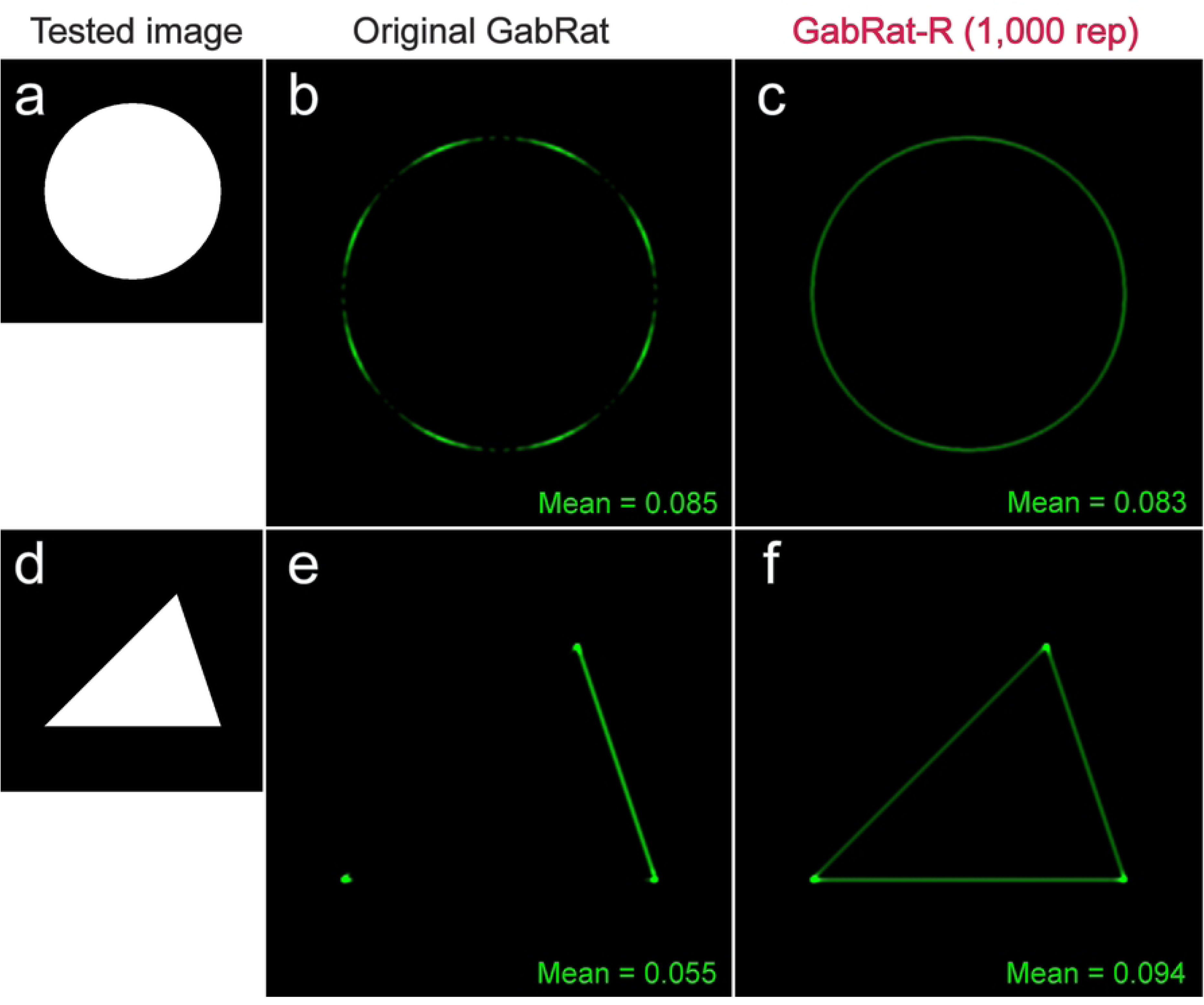
Comparison of the original GabRat and GabRat-R. (a) A tested image. This is the simplest pattern, in which a white circular object is put on the black background. Since there is apparently no ’false-edge’ in this picture, the edge disruption effect is expected to be constant and nearly zero everywhere on the edge points. (b-c) Results of GabRat and GabRat-R (1,000 iterations), respectively. Intensity of the green signals represents GabRat values (0.0 to 1.0) at each location. An angle-dependent, periodic pattern is seen in the original GabRat. (d-f) Similar comparison using a triangular shape. *Sigma* = 6, *Fx* = 2, *nAngles* = 4.

**Fig. 5.**
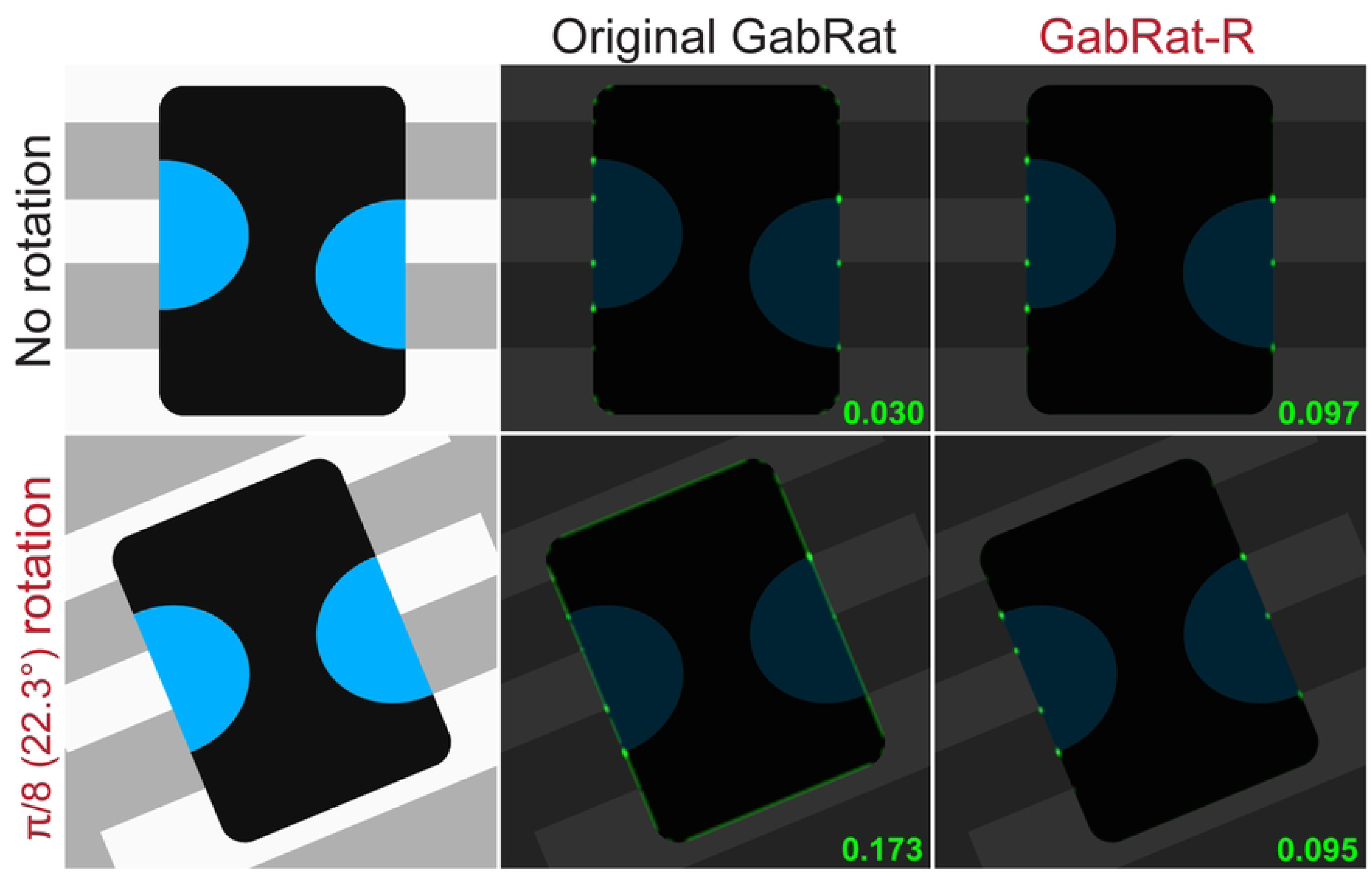
Rotation-dependent false signals in the original GabRat and averaged false signals in GabRat-R. *GabRat* values at each location are presented as the green signals superimposed on the original images. The mean values are shown at the bottom-left of the panels.

**Table 1.**
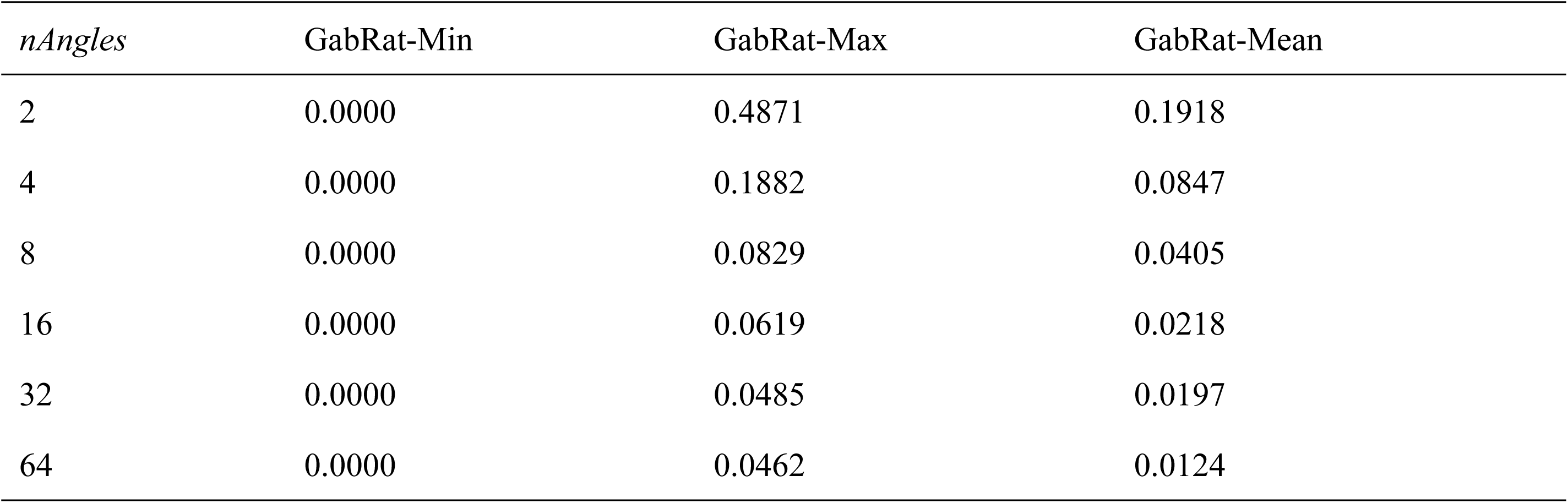
Effect of the number of Gabor filters (*nAngles*) on GabRat calculation. Target object is a white circle (*R* = 200, *EdgeCount* = 1132) on the black background. *Sigma* = 6.0, *Fx* = 2.

Why does such false signal occur in GabRat calculation? The answer is because there is a mismatch between the discrete edge angle (*θ_par_*) determined by the limited number (*nAngles*) of Gabor filters and the actual angle of the edge outline, as mentioned in the previous section. Fig 6 shows the same circular model image as Fig 4a, in which the binary image (i.e. a temporal image that GabRat program automatically generates, see Fig 3a) is identical to the real image. At the upper-most position of the circle (Fig 6, position I), the parallel angle *θ_par_* = 0° and *E_par_* takes a large negative value (since the kernel filter pattern is opposite to the black-and-white pattern of the local image) whereas *θ_ort_* = 90° and *E_ort_* = 0 (since the kernel filter pattern is perpendicular to the local image pattern); therefore, *GabRat* = |*E_ort_*| / (|*E_par_*| + |*E_ort_*|) = 0. However, at the second position (Fig 6, position II), *θ_par_*and *θ_ort_* remain unchanged (*θ_par_* = 0°, *θ_ort_*= 90°) but *E_ort_* takes a nonzero value; therefore, *GabRat* > 0. The easiest solution to reduce the false signal is to increase the *nAngles* parameter, since *θ_par_* becomes closer to the actual edge angle (Table 1). However, increasing *nAngles* parameter significantly decreases the mean GabRat values (Table 1), therefore, GabRat values calculated by different *nAngles* parameter cannot be directly compared with those determined in the previous studies, which typically use *nAngles* = 4.

**Fig. 6.**
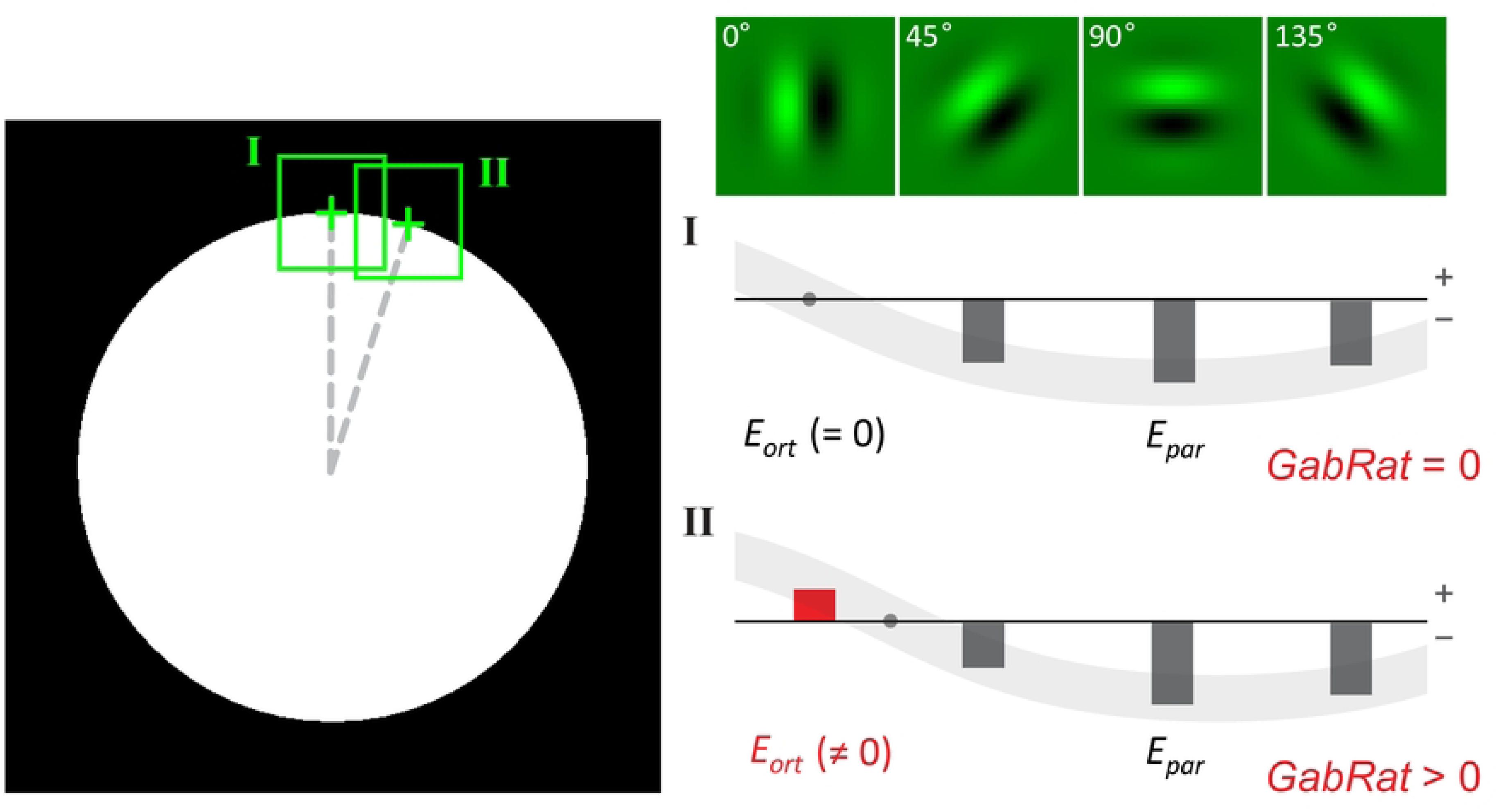
The cause of the angle-dependent false signals in the original GabRat calculation. The model image and GabRat parameters are the same as Fig. 5a. Since the binary image is identical to the real image, *E_k_* is always equal to *E’_k_*. (a) At the upper-most position of the circle edge, *E_par_* takes a large negative value (because the black-and-white pattern of the local area is opposite to that of the kernel filter) whereas *E_ort_* = 0, thus, *GabRat* = 0. (b) However, at the second position, *E_ort_*takes a small negative value due to the discrepancy between actual tangential angle and the kernel filter angle. As a result, *GabRat* takes a nonzero value.

### GabRat-R

Here we developed an improved method, namely GabRat-R, which can almost flatten the false signal but still keeps a sufficient mean-level compatibility with the original GabRat method. GabRat-R essencially uses the same number of Gabor filters (typically, *nAngles* = 4) to determine *θ_par_* as used in the original GabRat. Further, GabRat-R internally uses an additional *baseAngle* parameter, which represents an offset rotation angle of each Gabor filter (S1c Fig). The GabRat-R repeatedly calculates the GabRat values (e.g. 1,000 repetitions) with a randomly generated *baseAngle* parameter (0 ≤ *baseAngle* < 180°) based on Mersenne twister algorithm [21] for each iteration. The number of iterations is specified by the *nRepeat* parameter, which is typically set to 1,000.

In the cases of Fig 4, GabRat-R nearly flattened the false signal throughout the edge outline of the object, but the mean GabRat values were still close to that of the original GabRat method (Fig 4c, f). Another advantage of GabRat-R over GabRat is that the flattened false signals makes the spatial structure of the edge disruption intensity more obvious. Fig 7 illustrates the difference in the spatial structures of GabRat vales between the original GabRat and GabRat-R. In this verification experiment, a circular white model object with disruptive black stripes was put on a background image with grey tiled patterns (Fig 7a). There were four types of edge points in terms of disruptive patterns: α) the points where a boundary of the stripe pattern of the object (black & white) intercepts with the edge outline of the object, β) the points where a boundary of the stripe pattern of the background (dark grey & grey) intercepts the edge outline of the object, α+β) both α and β happen coincidently, and γ) the other locations where no disruptive pattern exists. Since the contrast of the object’s coloration was higher than that of the background pattern, it was expected that there are stronger GabRat signals at α locations and weaker signals β locations. In the original GabRat calculation, some β signals nearly disappeared due to the interference of background noise, whereas those local GabRat peaks were clearly seen in the result of GabRat-R (Fig 7b, c). These α and β peaks were also clearer in the histogram of GabRat values in GabRat-R calculation (Fig 7d, e). The difference in signal intensities between the original GabRat and GabRat-R at the same location tended to be larger when the GabRat value shows intermediate edge disruption (GabRat ≈ 0.5) (Fig 7f).

**Fig. 7.**
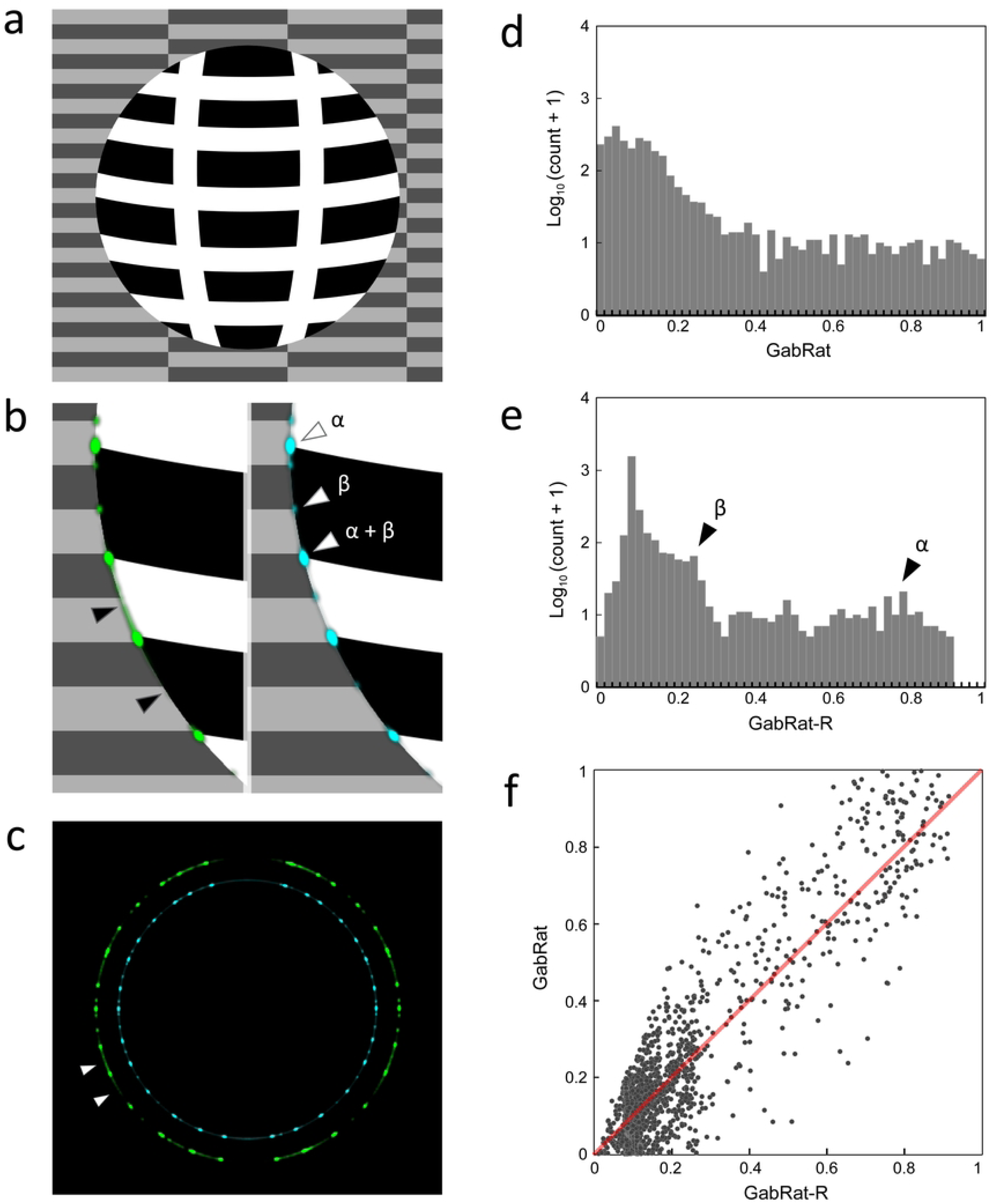
Difference of spatial structures of GabRat signals in the original GabRat and GabRat-R. (a) A model object put on a background image. The model object has stripe patterns with high contrast whereas the background consists of block-like patterns with low contrast. (b, c) Difference in fine signal distributions between GabRat (green) and GabRat-R (blue). Arrowheads indicate (α) the location where high contrast patterns on the object intersects the edge outline and (β) the location where low contrast patterns on the background intersects the edge outline. (d, e) Frequency distributions of the signal intensity of GabRat and GabRat-R, respectively. (f) Correlation between GabRat and GabRat-R. A dot represents the intensities of GabRat and GabRat-R at a specific pixel on the edge outline. A red line indicates the equilibrium line (*y* = *x*).

In addition to the new random base angle feature, the algorithm of GabRat itself has also been improved to significantly increase the calculation speed, which is important for the iterative calculation of GabRat in the GabRat-R method. Our new GabRat program is written in C++ and runs for native CPUs, giving it a speed advantage over the original GabRat program that has been developed as a Java plugin function, micaToolbox [15] of Image J software [19][20] and runs in the Java virtual machine. Moreover, GabRat-R uses the multithreading technology to run each iterative GabRat calculation in parallel, making its computational speed faster again. To evaluate the efficiency of algorithm besides the difference in the running environment, we first converted the Java program of original GabRat into C++ program mostly as it was (in word-by-word manner, since these two computer languages have quite similar syntax), and composed it into a single C++ function. For the detailed process of conversion, see the supplementary information. After that, two C++ source codes of the original GabRat and our new GabRat program were compiled in Visual C++ 2005 (Microsoft) with the moderate optimization for execution speed (/O2). As a test image, we prepared a composite image of a flightless Easter Egg weevil, *Pachyrynchus tobafolius* (Coleoptera: Curculionidae), placed on a natural background (S2 Fig). To measure the computational speed of the two programs, they were run for a single iteration and 1,000 iterations with common conditions: *sigma* = 5.0, *gamma* = 1.0 and *Fx* = 2.0. As a result, the improved GabRat program achieved approximately three times speedup compared to the original GabRat (Table 2). GabRat-R can run the improved GabRat program in parallel and significantly improved the computational speed again when the multithreading technology was available (Table 2).

**Table 2.** Comparison of the calculation time among the original GabRat implementation (rewritten in C++), the improved algorithm of our GabRat, and the GabRat-R which computes the improved GabRat in parallel.

| Sigma | Iteration | Original GabRat | Improved GabRat | GabRat-R |  |  |  |  |  |
| --- | --- | --- | --- | --- | --- | --- | --- | --- | --- |
|  |  |  |  | 1 thread | 2 threads | 4 threads | 6 threads* | 8 threads | 12 threads |
| 4 | 1 | 46 ms | 16 ms | - | - | - | - | - | - |
|  | 1,000 | 39.1 s | 15.6 s | 12.2 s | 6.3 s | 3.2 s | 2.2 s | 2.5 s | 2.3 s |
| 8 | 1 | 125 ms | 47 ms | - | - | - | - | - | - |
|  | 1,000 | 124.7 s | 49.5 s | 45.7 s | 23.0 s | 11.9 s | 8.8 s | 9.5 s | 8.5 s |

\* Default number of threads in the tested environment (Intel Core i7-990X, 6 cores/12 threads, overclocked to 4.5 GHz). The GabRat-R program automatically set it to the number of CPU cores.

### GabRat-RR

The original GabRat method first requires a composite image where a target object is placed on a particular background image (e.g. Fig 7a). GabRat calculates the edge energies by looking at patterns in both the object and the background near the object’s edge, and thus, the background essentially affects the GabRat value (e.g. Fig 7b). Natural backgrounds, such as tree leaves and tree trunks, usually consist of different local patterns and often have angle-dependent components. Therefore, intensity of the edge-disrupting effect may also depend on the position and the angle where the foreground object is placed.

GabRat-RR method was thereby developed to average the effects of the heterogeneity and anisotropy of background images. Prior to the GabRat-RR computation in *Natsumushi* 2.0 software, a foreground image (i.e. a picture containing a target object) (Fig 8a) and a background image must be prepared separately. To specify the range of the target object (e.g. the body part of a weevil excluding antennae and legs), a region-of-interest (ROI) must be specified on the foreground image (Fig 8b). GabRat-RR repeatedly generates composite images of the foreground object placed against the background image, using random positions and rotation angles (Fig 8c, d) and calculates the GabRat values. In each iteration, GabRat-RR also generates a random *baseAngle* parameter (as mentioned in GabRat-R) to rotate the Gabor filters and cancel the angle-dependent false signals. This double randomization feature is the origin of the name GabRat-RR. The number of the iteration is specified by *nIndividual* parameter. As a result, each location on the object’s contour has *nIndividual* distinct GabRat values because the adjacent background patterns are different. At the end of the iterated computation, GabRat-RR summarizes the results in two ways: 1) list of the mean GabRat values for each iteration, in which each value represents the intensity of edge disruption of an entire shape, and 2) means of the GabRat values at each location of the object’s contour. The first summary is related to the heterogeneity and anisotropy of the background, and the second summary shows the spatial distribution of GabRat intensity on the object.

**Fig. 8.**
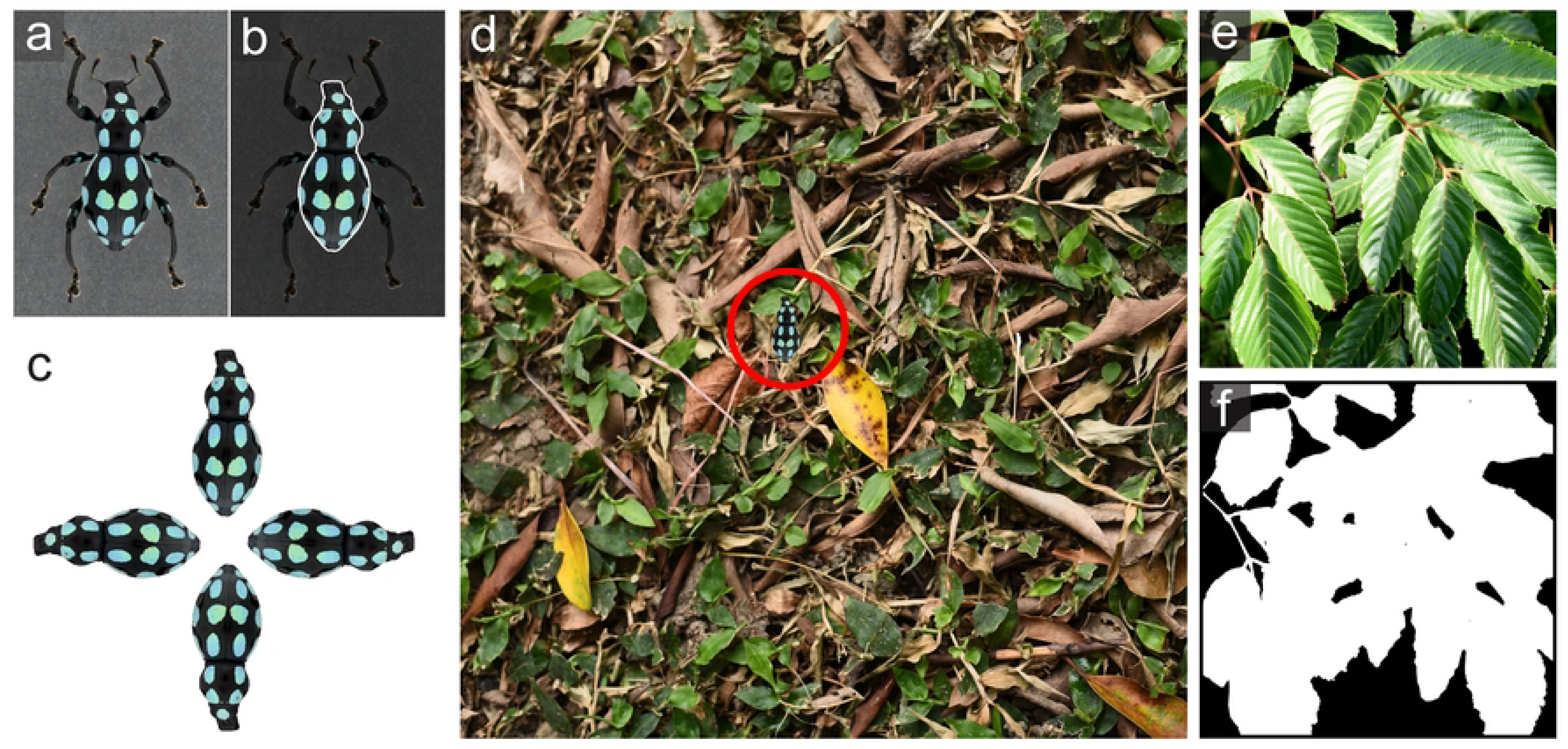
An analytical procedure of GabRat-RR using Natsumushi 2.0. (a) An original insect image. (b) A region of interest (ROI) on the insect image. (c) Four rotated insect images by π/2 step. (d) GabRat-RR randomly selects one of the rotated ROIs, puts it at the random position on the background, and calculates a *GabRat* value with using a randomly generated *baseAngle* parameter. This procedure is repeated *nIndivudual* times. (e-f) Mask images to specify the area where the object is allowed to put.

Rotation angle of the target object is currently restricted in four discrete angles: 0°, 45° (1/2π), 90° (π), and 135° (3/2π) (Fig. 8c). That is because other intermediate rotations always cause pixel-level rearrangement of the object’s contour and change the number of edge pixels, which makes it difficult to summarize the results in the second form. We have already found a technical solution that allows the use of continuous random rotation angles, and this improvement will be applied in future versions of the GabRat-RR program.

Natural background images often contain areas which are not suitable for the target object to be placed. For example, a flightless *Pachyrynchus* weevil cannot stay in the empty space between leaves (Fig. 8e). In such cases, GabRat-RR can use an optional mask (black and white) image to specify the valid areas of background (Fig. 8f). If a mask image is present, GabRat-RR program will place the target object only in the valid areas specified by white pixels in the mask image.

### Comparison of original GabRat, GabRat-R and GabRat-RR

This study demonstrated that GabRat value from the original GabRat method is significantly affected by the angles of edge outline, especially when it includes straight lines (Figs 4, 5). Most previous studies used isosceles triangles as template shapes, mimicking a moth perched on an environmental background [22][23]. In such studies, false GabRat signals might possibly have happened depending on two factors: 1) the shape of the triangle, and 2) the direction of head. Here, we aimed to verify the existence of biased GabRat evaluation in those common experimental cases. For a model organism, we choose a moth species, *Dinumma deponens*, which is common in Taiwan and has typical disruptive colorations on the wing (Fig 9a). A living specimen of *D. deponens* was photographed in Ma-Mei forest trail, Hsinchu County, Taiwan (Fig 9a), and the outlines of the moth was transformed into isosceles triangular shapes with vertex angles of 90° (right triangle: Fig 9b), 60° (equilateral triangle: Fig 9c) and 45° (acute triangle: Fig 9d) by the thin plate spline (TPS) deformation using *Natsumushi* 2.0 software [24][25]. The deformed images were trimmed by exact isosceles triangles, resized so that the area of the triangles become approximately 10,000 pixels, and converted into grayscale images (Fig 9b-d). For background images, two pictures of the leaves of *Bischofia javanica* and *Macaranga tanarius* trees and two pictures of the trunks of *B. javanica* in different appearances were resized into 3,000 × 3,000 pixels so that the length of each side corresponds to 15 cm in the real objects, and finally converted into grayscale images (Fig 9e-h).

**Fig. 9.**
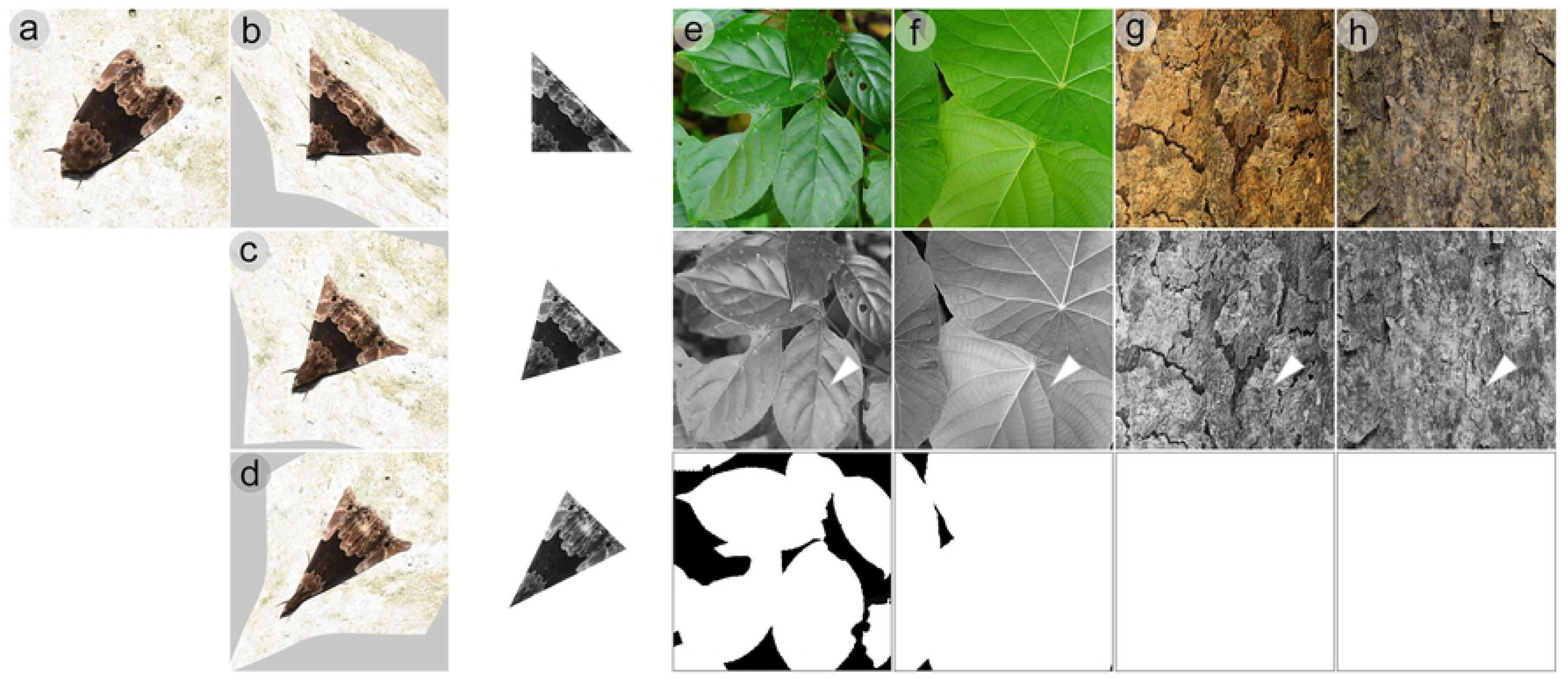
A natural model images and natural backgrounds used for the comparison of the original GabRat, GabRat-R and GabRat-RR. (a) An image of the moth *Dinumma deponens.* (b-d) To simulate the experimental conditions of previous studies, the moth image was deformed into three different triangular shapes using TPS transformation. The deformed images were then cut into acculate triangles, resized so that they all have the same area size, and converted into gray images. (e-h) Background pictures were also converted into gray images. Arrowheads indicates the fixed location where the triangular moth images were put in GabRat-R analysis. For GabRat-RR, mask images were used to specify the valid area ranges where the moth images were allowed to put.

To test the difference in original GabRat and the GabRat-R, those three triangular moth images were put on a fixed position of each background pictures (arrowheads in Fig 9e-h) with no rotation (0°) andπ/8 (22.5°) rotation. The common parameters for the original GabRat and GabRat-R were: *Sigma* = 6, *Gamma* = 1, *Fx* = 2, and *Baseangle* = 0. For the original GabRat, the ‘random base angle’ function was disabled and *nRepeat* = 0 (that enforces the application to use the original GabRat method), whereas GabRat-R uses the ‘random base angle’ function with *nRepeat* = 1,000. The result showed that the difference in GabRat-R and GabRat-RR tend to be greater when using a right triangle shape, especially when the moth images were rotated by 22.5° (Fig 10a). Additionally, we also tested GabRat-RR to clarify the effect of heterogeneity in background. Since the pictures of the tree leaves contain some empty areas where a moth cannot stop, we use the mask images to specify the valid region (Fig 9e, f). When the results of GabRat-RR were compared with those of GabRat-R at two rotations, substantial differences were found within each combination of moth shapes and backgrounds (Fig 10b). However, no clear tendency was observed between those two analyses, perhaps because of the low heterogeneity of the background images.

**Fig. 10.**
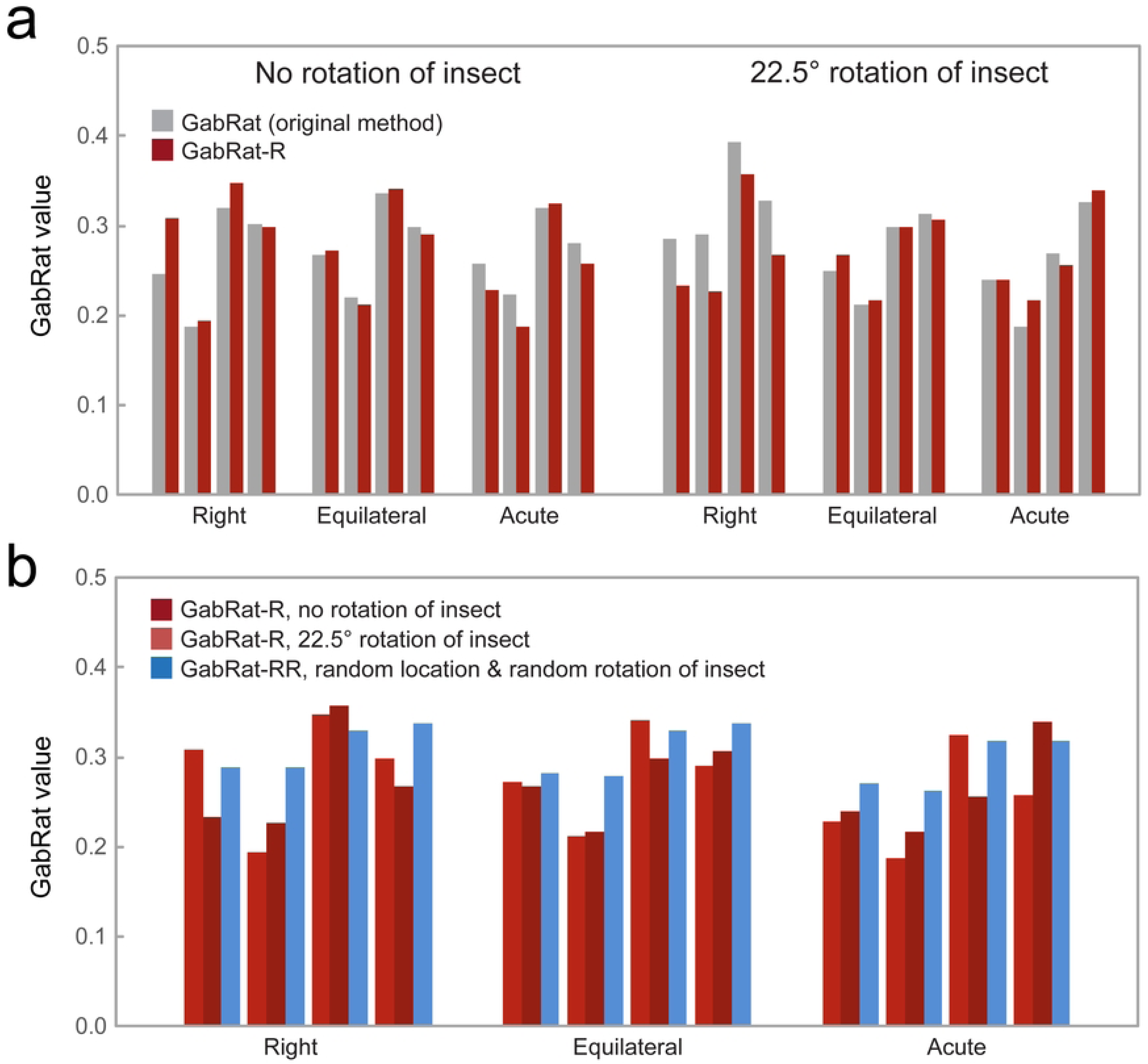
Comparison of different GabRat methods. (a) Comparison of the original GabRat and GabRat-R, using the moth images of three triangular shapes putting at the fixed position of the four different natural backgrounds (see Fig. 9). (b) Comparison of GabRat-R (fixed position and rotation angle) and GabRat-RR (randomized position and rotation angle, *nIndividual* = 1,000).

## Conclusion and Applications

The original GabRat calculation has an intrinsic issue that generates false signals and they may affect the mean *GabRat* value in certain cases, especially when the target shapes have long, straight edge outline s with intermediate angles. Our GabRat-R can average such angle-dependent issues and provide unbiased comparisons among different shapes. Furthermore, GabRat-RR is useful to account for the heterogeneity and anisotropy of background images.

## Program Availability

All the image operations, analytical features and visualization tools used in this study (i.e. GabRat-R/RR, ROI handling, color conversions and TPS deformation) are implemented in *Natsumushi* 2.0 software [24][25], which is available from the author’s website: https://sites.google.com/site/mtahashilucanid/program/natsumushi.

## Acknowledgements

We greatly thank Chun-Yu Lin (National Taiwan Normal University) for providing insect photographs.

## Author contributions

MT and ML prepared the materials. MT designed and wrote the computer programs. MT and ML conducted the experiments and created the figures. All authors wrote and reviewed the manuscript.

## Additional Information

Supporting information accompanies with this paper.

## Supporting information

**S1 Fig. Changes of the Gabor kernel filter.**

**S2 Fig. Images used to test the computational speed of different GabRat programs.**

**S3 File. Additional methods**

